# Microbial mutualism generates multistable and oscillatory growth dynamics

**DOI:** 10.1101/2022.04.19.488807

**Authors:** Daniel B. Amchin, Alejandro Martínez-Calvo, Sujit S. Datta

**Affiliations:** Department of Chemical and Biological Engineering, Princeton University, Princeton, NJ 08544, USA; Princeton Center for Theoretical Science, Princeton University, Princeton, NJ 08544, USA

## Abstract

Microbial communities typically comprise multiple different species with an intricate network of interactions, ranging from competitive to cooperative, between them. How does the nature of these inter-species interactions impact overall community behavior? While the influence of purely competitive interactions is well-studied, the opposite case of mutualistic interactions—which are also prevalent in many naturally-occurring communities—is poorly understood. Here, we address this gap in knowledge by mathematically modeling a well-mixed two-species community of aerobes and anaerobes having mutualistic metabolic interactions between them. Despite the simplicity of the model, we find that it reproduces three characteristic experimental findings. In particular, in response to changes in the fluxes of exogenously-supplied carbon and oxygen, the community adopts two *distinct stable states* with differing fractions of aerobes and anaerobes. These states are *bistable*, capable of arising under identical environmental conditions; transitions between the two are therefore history-dependent and can give rise to *oscillations* in the bacterial and chemical concentrations. Moreover, using the model, we establish biophysical principles describing how oxygen depletion and nutrient sharing jointly dictate the characteristics of the different states as well as the transitions between them. Altogether, this work thus helps disentangle and highlight the pivotal role of mutualism in governing the overall stability and functioning of microbial communities. Moreover, our model provides a foundation for future studies of more complex communities that play important roles in agriculture, environment, industry, and medicine.

## I. INTRODUCTION

Microbial communities often comprise different species having distinct functions, metabolic capabilities, and requirements for survival; nevertheless, they can stably coexist as a group, often in dynamic environments with strongly-fluctuating nutrient availability. How is this coexistence achieved?

As a necessary first step toward addressing this question, numerous studies have documented the varying ways in which different species in a community interact, ranging from mutually-harmful competition to mutually-beneficial cooperation. This network of interactions can give rise to fascinating emergent behaviors whose occurrence is remarkably consistent across diverse communities. For example, a common finding is that microbial communities can have *multiple stable states*, each characterized by its own unique composition of the different coexisting species, and each of which is stable under different environmental conditions^1–15^. In some cases, these states are *multistable*—i.e., multiple stable states can arise under identical conditions—leading, for example, to *hysteretic* behavior in which the state of the community depends not just on current conditions, but also on the history of how they were established^2–4,9,16–26^. Multistability can also lead to dynamic behavior in which the community continually fluctuates between multiple states, often with periodic *oscillation*^8,13,15,19,27–33^.

Competition for limited resources is typically inherent in multi-species communities; hence, much work has focused on understanding ways in which such behaviors can emerge in communities with purely competitive interactions^13,18,19,23–25,28,34–38^. However, a growing body of research is revealing that mutualistic interactions, such as cross-feeding of metabolites, also arise and play critical roles in many naturally-occurring microbial communities^7,11,39–64^. For example, microbial mutualism regulates the consumption of marine particulate organic matter^4,5,65,66^, plant growth^67–69^, how carbon and nitrogen are fixed in or released from the ground beneath our feet^70^, and degradation of environmental contaminants^71^—with profound implications for biogeochemical processes in the world around us. Such interactions also play key roles in our own bodies. A prominent example is that of microbial communities in the gut, lung, and mouth, in which anaerobic bacteria can ferment large carbon-rich macromolecules that are inaccessible to nearby aerobes, releasing smaller byproducts that support the growth of the aerobes, which in turn help support anaerobic growth by consuming oxygen^2,72–75^—and collectively, such aerobe-anaerobe communities help maintain host health^76^, or conversely, contribute to disease^72,77–81^. Similar mutualistic aerobe-anaerobe communities also arise and play crucial roles in many other ecological and biotechnological settings^70,82–88^. Understanding how mutualism influences the overall behavior of a microbial community is therefore both fundamentally interesting and practically important for predicting and controlling a variety of environmental, agricultural, biomedical, and industrial processes.

Given that mutualistic interactions are prevalent in microbial communities, and that such communities frequently exhibit multistability, hysteresis, and time-dependent behaviors, we asked the questions: Can mutualism generate these complex behaviors? And if so, are there simple biophysical principles that describe these behaviors and the conditions under which they arise? Prior experiments on model two-species communities of aerobes and anaerobes in bioreactors^2,3,27,89^ provide useful guidance in addressing these questions. In particular, this prior work demonstrated experimentally that such communities can indeed exhibit multistability, hysteresis, and time-dependent behaviors. Moreover, it showed that many of these behaviors can be recapitulated using sophisticated models of the intricate network of metabolic interactions between cells, as reconstructed from genomic data^2,90–93^. However, such networks are made up of a multitude of vastly-differing interactions, ranging from competitive to cooperative; thus, the role played by mutualism in generating these behaviors is obfuscated, making the formulation of simple overarching biophysical principles challenging.

Here, we address this challenge by mathematically modeling a two-species community of aerobes and anaerobes having a simplified set of mutualistic interactions between them. Specifically, inspired by the prior studies noted above, we consider the case in which the anaerobes break down nonmetabolized complex carbohydrate to simple sugar that is shared by the entire community—but only under low-oxygen conditions that are established through aerobic consumption of oxygen. Remarkably, even in this highly-simplified community, we find that multistability, hysteresis, and time-dependent behaviors arise, mediated by carbon and oxygen fluxes just as in experiments. Moreover, the simplicity of our model enables us to distill out biophysical principles that quantify how oxygen depletion and nutrient sharing jointly enable coexistence—highlighting the pivotal role of mutualism in enabling coexistence without needing to invoke antagonism. These principles quantitatively capture the conditions under which different community behaviors arise in our model, providing a foundation for future studies of more complex multi-species microbial communities in a broad range of settings.

## II. METHODS

### A. Development of the governing equations

As an illustrative and well-characterized^2,3,27,89^ example, we consider the model two-species microbial community schematized in Fig. 1a(ii). This community is a mixture of aerobes (red in Fig. 1) and anaerobes (dark blue), whose metabolism and growth are either promoted or instead suppressed in oxygenated environments, respectively. The bacteria are described by a number concentration *b* with subscripts aer and an for aerobes and anaerobes, respectively. Genome sequencing and metabolic profiling of such communities indicate that the consumption and secretion of only a small number of metabolites often drives experimental outcomes^3–5,7,13,94^—providing a clue that the full network of metabolic interactions could be dramatically simplified while still generating realistic community behaviors. Hence, inspired by^2^, we focus on the case in which the anaerobes take up an exogenously-supplied complex carbohydrate (teal) that cannot be accessed by the aerobes, breaking it down into simple sugar molecules (green) that they either directly consume for their growth or liberate to be consumed by the entire microbial community; for simplicity, we do not consider any other compounds, such as short-chain fatty acids, that may also liberated upon carbohydrate breakdown. The aerobes consume oxygen (magenta)—thereby providing favorable conditions for the anaerobes—while also consuming liberated simple sugar to utilize for their own growth. To examine the complex phenomena that can emerge in this model community, and to develop biophysical principles that can describe these phenomena, here we mathematically describe these mutualistic interactions by building on the framework of consumer-resource models commonly used in ecology^11,38,55,94^.

**Figure 1.**
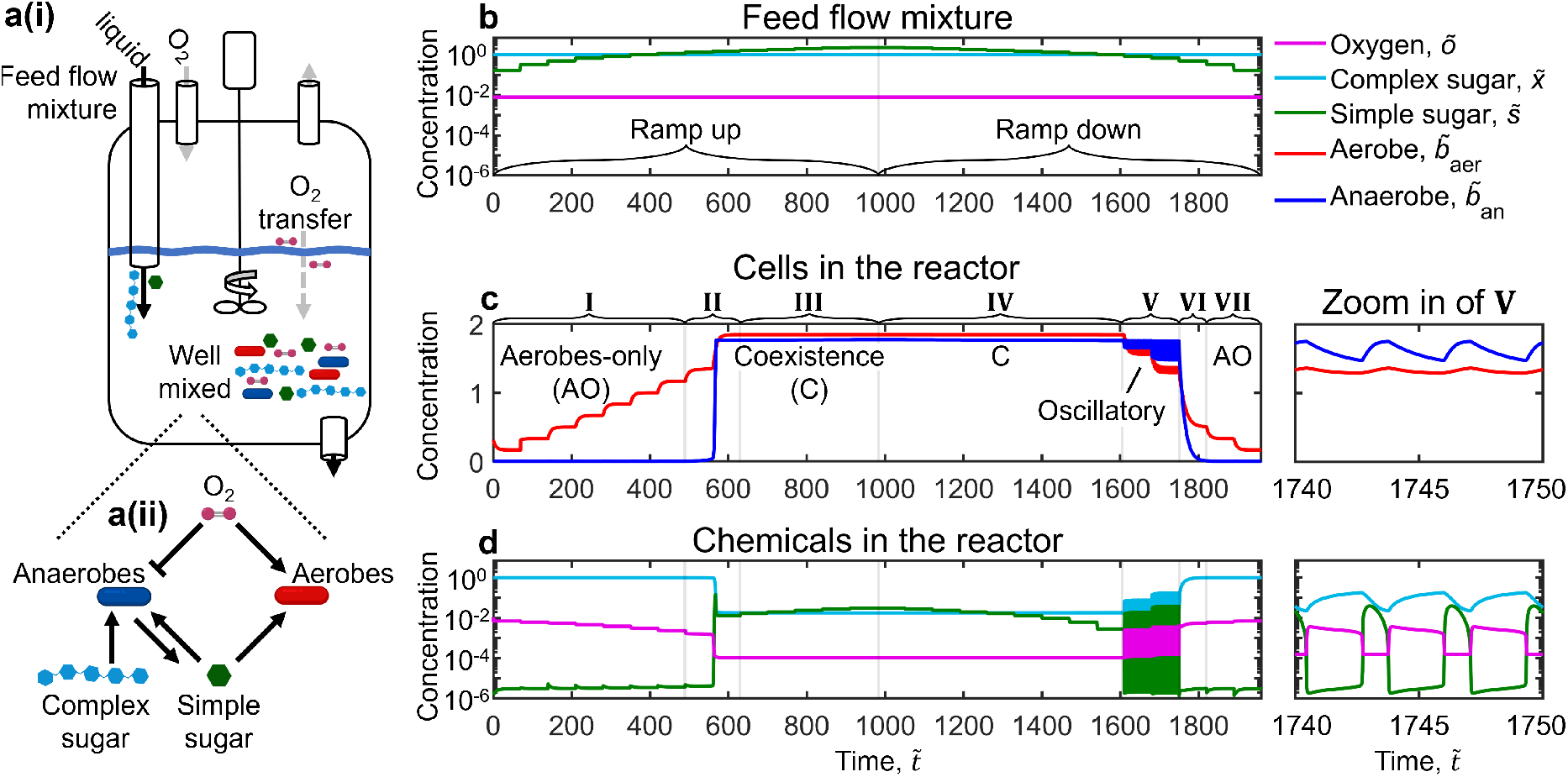
Model community of aerobes and anaerobes with mutualistic metabolic interactions exhibits two stable states, with bistable and oscillatory transitions between them driven by changing nutrient and oxygen fluxes. **a(i)** We consider a continuously-stirred tank reactor containing a well-mixed liquid culture of aerobes, anaerobes, dissolved oxygen, complex carbohydrate (labeled as complex sugar for brevity), and simple sugar. A liquid feed flow enters the reactor with dissolved complex carbohydrate and simple sugar, along with oxygen gas maintained at a constant partial pressure in the head space above the liquid. The rates of liquid outflow and inflow are matched, maintaining a constant liquid volume in the reactor. **a(ii)** Schematic of the mutualistic interactions between bacteria. Anaerobes take up complex carbohydrate and degrade it to simple sugar, some fraction of which is utilized solely by anaerobes, and the remainder of which is liberated to be shared by both anaerobes and aerobes for their growth. Aerobes consume oxygen, which inhibits (enhances) anaerobe (aerobe) activity. **b–d** show the results of an exemplary numerical simulation of our model with intermediate nutrient sharing (Ξ = 0.5) and oxygen abundance *δ* = 3.9 × 10^−3^, using parameter values that are representative of experiments as given in Table II. The composition of the incoming feed is given in **b**; to examine the influence of changing carbon fluxes, we progressively increase (ramp up) and then decrease (ramp down) the concentration of simple sugar in the feed. The resulting changes on the concentration of cells and chemicals in the reactor are given in **c** and **d**, respectively. Initially, the community is in the aerobes-only (AO) state (I), then abruptly transitions (II) to a state in which aerobes and anaerobes coexist (C, III–VI). Upon ramping down the simple sugar feed concentration, the C state persists, indicating hysteresis in the transitions between states. Further decreasing the simple sugar feed concentration gives rise to oscillations in the bacterial and chemical concentrations (magnified views to the right) before the community ultimately transitions back to the AO state (VII). All quantities are given in dimensionless form indicated by tildes; we rescale {*t, x, s, o, b*_an_, *b*_aer_} by the characteristic quantities 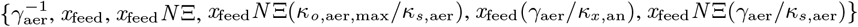 as detailed in the text.

**Figure 2.**
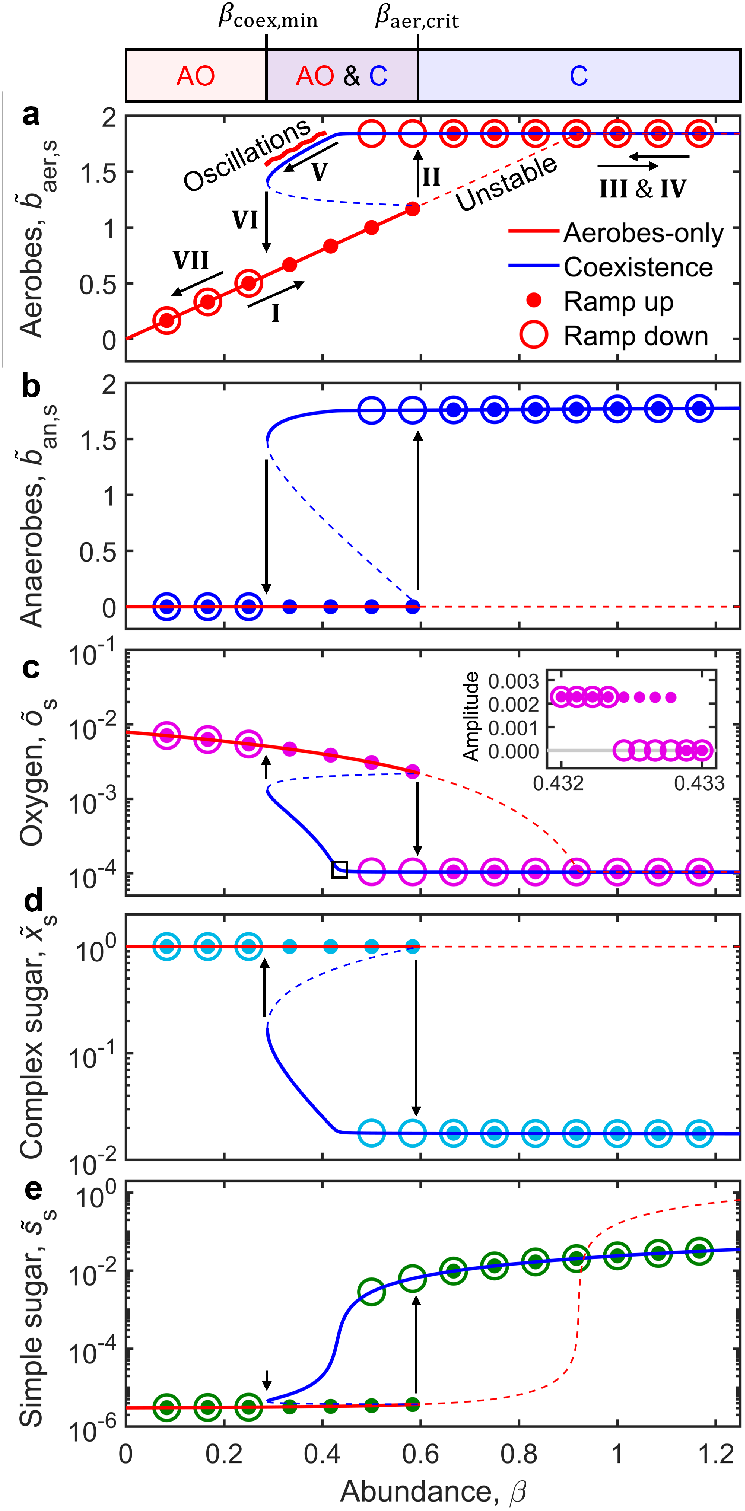
Phase space of microbial steady states. Panels **a–b** show the steady-state aerobe and anaerobe concentrations at each value of the simple sugar abundance *β* for the simulation shown in Fig. 1; the corresponding steady-state chemical concentrations are given in panels **c–e**. Filled and open circles show ramp up (increasing *β*) and ramp down (decreasing *β*), respectively. Red and blue curves show the analytical solutions predicted by our theory (Eqs. 11–15); solid and dashed lines indicate the stable and unstable states, respectively. Initially the microbial community is in the aerobes-only (AO) state (I), and then transitions (II) to the coexistence (C) state (III) above *β* = *β*_aer,crit_ ≈ 0.6. Upon ramping down, the community persists in the C state (IV) over a broader range of *β*, indicating hysteretic behavior. It eventually exhibits oscillations (V) and then transitions (VI) back to the AO state (VII) below *β* = *β*_coex,min_ ≈ 0.28. Inset to **c** shows the amplitude of oscillations in oxygen concentration just near the onset of oscillatory dynamics, in the narrow region indicated by the small black box in the main panel; the oscillation amplitude displays a hysteresis loop characteristic of a subcritical Hopf bifurcation, with the grey line indicating zero amplitude for clarity. The overlying bar indicates monostable AO, bistable AO and C, and monostable C regimes demarcated by the critical values *β*_coex,min_ and *β*_aer,crit_. Black arrows indicate the direction of transitions between states. All input parameters are the same as in Fig. 1 except 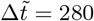, which is longer to ensure that simulations reach the long-time steady state. All quantities are given in dimensionless form indicated by tildes; we rescale {*x, s, o, b*_an_, *b*_aer_} by the characteristic quantities {*x*_feed_, *x*_feed_*N*Ξ, *x*_feed_*N*(*κ*_*o*,aer,max_/*κ*_*s*,aer_), *x*_feed_(*γ*_aer_/*κ*_*x*,an_), *x*_feed_*N*Ξ(*γ*_aer_/*κ*_*s*,aer_)} as detailed in the text.

**Figure 3.**
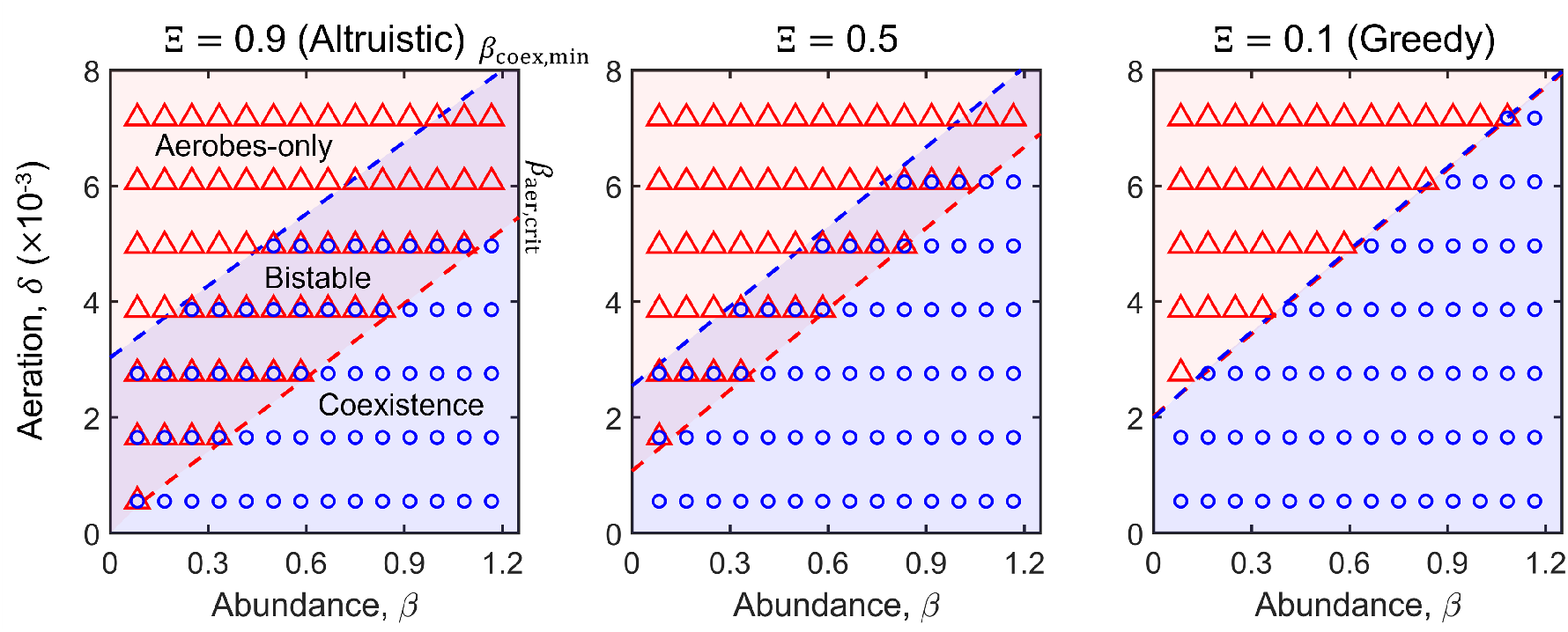
State diagrams for microbial communities under varying simple sugar and oxygen abundance. Symbols indicate the steady state reached by the microbial community in numerical simulations such as those in Fig. 2, holding a given value of the oxygen abundance parameter *δ* during ramp up (increasing simple sugar abundance *β* from 0 to 1.2) and ramp down (decreasing *β* from 1.2 to 0); red circles and blue triangles show the monostable aerobes-only (AO) and coexistence (C) states, respectively. Superimposed red circles and blue triangles show the bistable regime in which the AO state manifests during ramp up, while the C state manifests during ramp down. The transitions from AO to C and vice versa are described by the critical values *β* = *β*_aer,crit_ and *β* = *β*_coex,min_, respectively. The background shading and dashed lines show the stable states and critical *β* values predicted by our analytical steady-state solutions given by Eqs. 11–15. The analytical theory and numerical simulations show excellent agreement, except as expected for the case of large *δ*, for which *β*_aer,crit_ exceeds the upper limit of the *β* explored; thus, coexistence is never reached and the bistable regime is precluded. In all cases, coexistence requires both oxygen depletion and nutrient sharing (small *δ* and large *β*). Moreover, the regime of bistability over which coexistence persists narrows in going from the “altruistic” limit of strong nutrient sharing (left panel Ξ = 0.9) to the “greedy” limit of weak nutrient sharing (right panel, Ξ = 0.1) by anaerobes.

**Figure 4.**
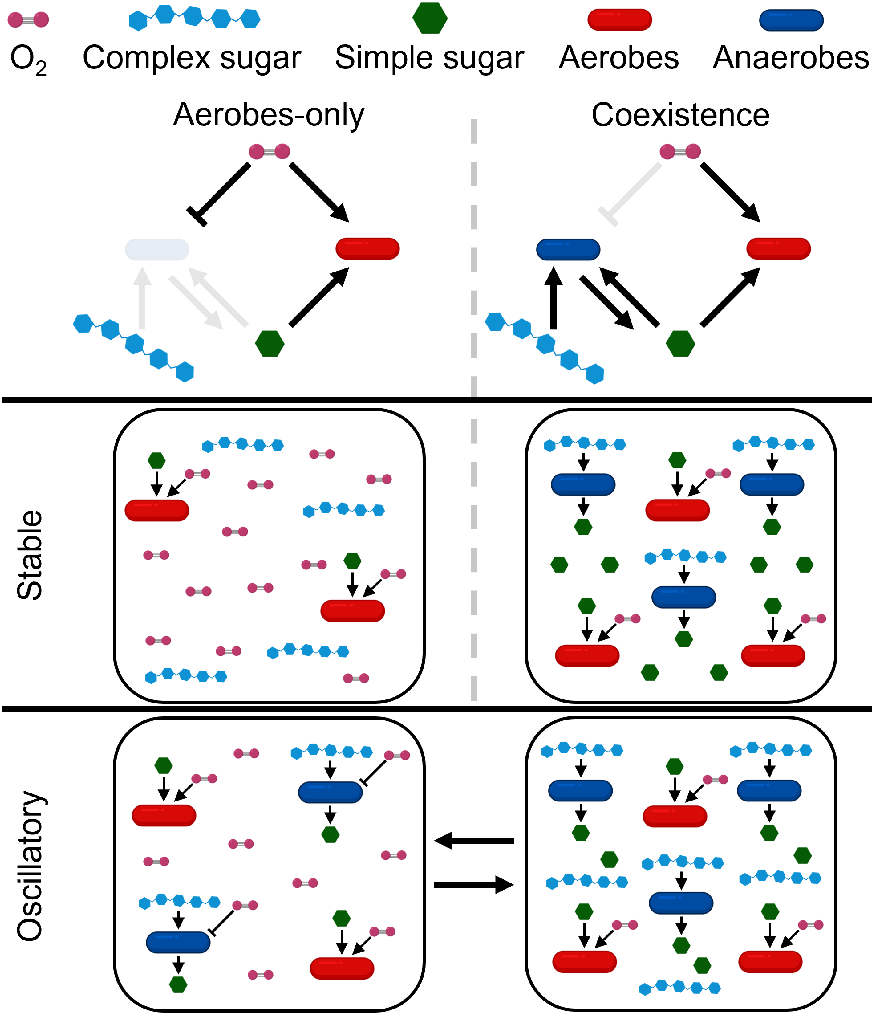
Summary of the different behaviors that arise in this microbial community. At low simple sugar abundance, the community is in the stable aerobes-only state (top and middle left) in which solely aerobes consume simple sugar, but are not concentrated enough to deplete oxygen for anaerobic growth. At high simple sugar abundance, the community transitions to the stable coexistence state (top and middle right) in which aerobes sufficiently deplete oxygen to sustain anaerobic growth, which in turn enables simple sugars to be liberated from complex carbohydrate and shared with the overall community. Upon subsequently decreasing the simple sugar abundance, before it transitions back to the aerobes-only stable state, the community oscillates between the low-oxygen (bottom right) and high-oxygen (bottom left) metastable states. In the former state, complex carbohydrate is rapidly broken down by anaerobes, causing aerobes to eventually run out of simple sugar and diminishing their ability to deplete oxygen—driving a transition to the latter state. In the latter, anaerobic breakdown of complex carbohydrate is slower, which enables complex carbohydrate and the subsequently liberated simple sugar to eventually become replenished, causing aerobes to increasingly deplete oxygen—driving a transition back to the former state.

#### Reactor configuration

In particular, we consider these coupled chemical and bacterial dynamics in the well-defined environment of a continuously-stirred tank reactor (CSTR)^2,27^, as shown in Fig. 1a(i); because all chemical and bacterial species are well-mixed, their concentrations are only functions of time, not space, in our model. In this configuration, a liquid feed flow containing dissolved complex carbohydrate at a concentration *x*_feed_ and simple sugar at *s*_feed_ enters the reactor at a fixed volumetric flow rate; in general, we use *x* and *s* to describe the number concentrations of complex carbohydrate and simple sugar in the reactor, respectively. The temperature and oxygen partial pressure in the head space of the reactor are held constant, thereby prescribing the saturation concentration of oxygen in the reactor liquid *o*_sat_; hence, the dissolved molecular oxygen concentration *o* changes at a rate *γ_o_*(*o*_sat_ – *o*), where *γ_o_* is the transfer rate coefficient^2^. The well-mixed liquid in the reactor exits at the same flow rate so as to maintain a constant internal volume. The flow rate divided by reactor volume then defines the dilution rate at which cells and carbon sources are refreshed to their feed flow quantities. We take *γ_d_* ≪ *γ_o_*; therefore, oxygen transfer is rapid and we do not include *γ_d_* in our equation describing oxygen dynamics. We also take the feed concentration of cells to be zero, and so, cells must grow within the reactor to avoid being diluted to a vanishing concentration and “washing out”.

#### Complex carbohydrate uptake

The anaerobes take up the exogenously-supplied complex carbohydrate at a rate *b*_an_*κ*_*x*,an_*g*_*x*,an_(*x*)[1 – *g*_*o*,an_(*o*)], where *κ*_*x*,an_ describes the maximum uptake rate per cell. In general, the Michaelis-Menten function *g*_*m*i_(*m*) ≡ *m*/(*m* + *m*_1/2,i_) quantifies the concentration dependence of the uptake of a substrate *m* ∈ {*o, x, s*} by species i ∈ {an, aer} relative to the characteristic concentration *m*_1/2,i_^95–101^. Thus, the uptake rate increases and eventually saturates with increasing concentration of the complex carbohydrate, mediated by the availability of anaerobic conditions at *o* ≲ *o*_1/2,an_^55^.

#### Consumption of simple sugar and oxygen

The anaerobes then break down the complex carbohydrate into simple sugar, a fraction Ξ of which are liberated by the anaerobes as a “common good” to be equally shared by all members of the microbial community, while the remaining 1 – Ξ are instead utilized solely by the anaerobes for their growth^2,11^. Thus, we describe the rate at which the anaerobes liberate simple sugar by *N*Ξ*b*_an_*κ*_*x*,an_*g*_*x*,an_(*x*)[1 – *g*_*o*,an_(*o*)], where the degree of polymerization *N* > 1 quantifies the number of simple sugar units liberated from a complex carbohydrate molecule. The anaerobes and aerobes then consume the simple sugar at total rates *b*_an_*κ*_*s*,an_*g*_*s*,an_(*s*)[1 – *g*_*o*,an_(*o*)] and *b*_aer_*κ*_*s*,aer_*g*_*s*,aer_(*s*)*g*_*o*,aer_(*o*), respectively; here, *κ*_*s*,an_ and *κ*_*s*,aer_ describe the maximum consumption rates per cell. Oxygen is concomitantly consumed by the aerobes at a rate *b*_aer_*κ*_*o*,aer_(*s*)*g*_*o*,aer_(*o*), where the consumption rate per cell depends on carbon availability^42,102^, and takes the form *κ*_*o*,aer_(*s*) ≡ *κ*_*o*,aer,min_ + (*κ*_*o*,aer,max_ – *κ*_*o*,aer,min_)*g*_*s*,aer_(*s*); this function interpolates between the basal consumption rate in starved conditions, *κ*_*o*,aer,min_, and the maximum consumption rate in sugar-replete conditions, *κ*_*o*,aer,max_. While we take *κ*_*o*,aer,min_ > 0 as is often the case, our results are unaltered if instead we take *κ*_*o*,aer,min_ = 0 (Fig. 5). Conversely, choosing *κ*_*o*,aer,min_ = *κ*_*o*,aer,max_, which completely removes the dependence of oxygen consumption by aerobes on simple sugar availability, influences some of the emergent community behavior (Fig. 6) as expected, as described further below.

**Figure 5.**
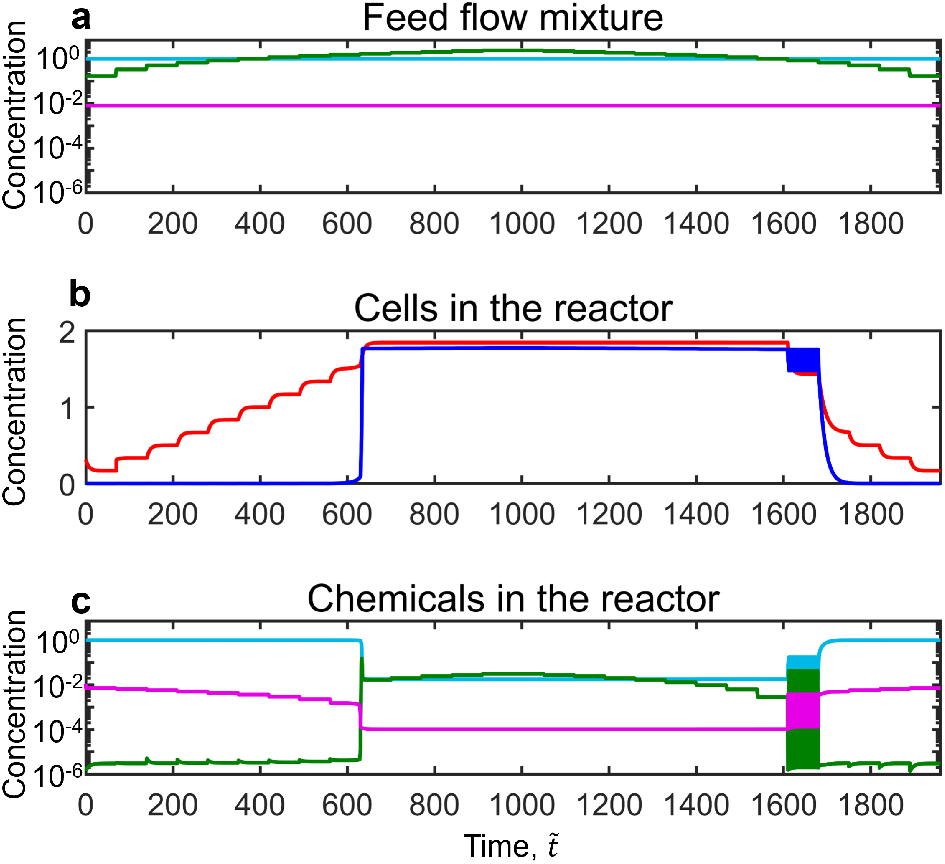
Our findings of bistability and oscillations are unchanged if the minimum oxygen consumption rate by aerobes is set to zero. While our main text results take *κ*_*o*,aer,min_ > 0 as is often the case, setting *κ*_*o*,aer,min_ = 0 yields nearly identical results, as shown in the Figure above, which corresponds to Fig. 1.

**Figure 6.**
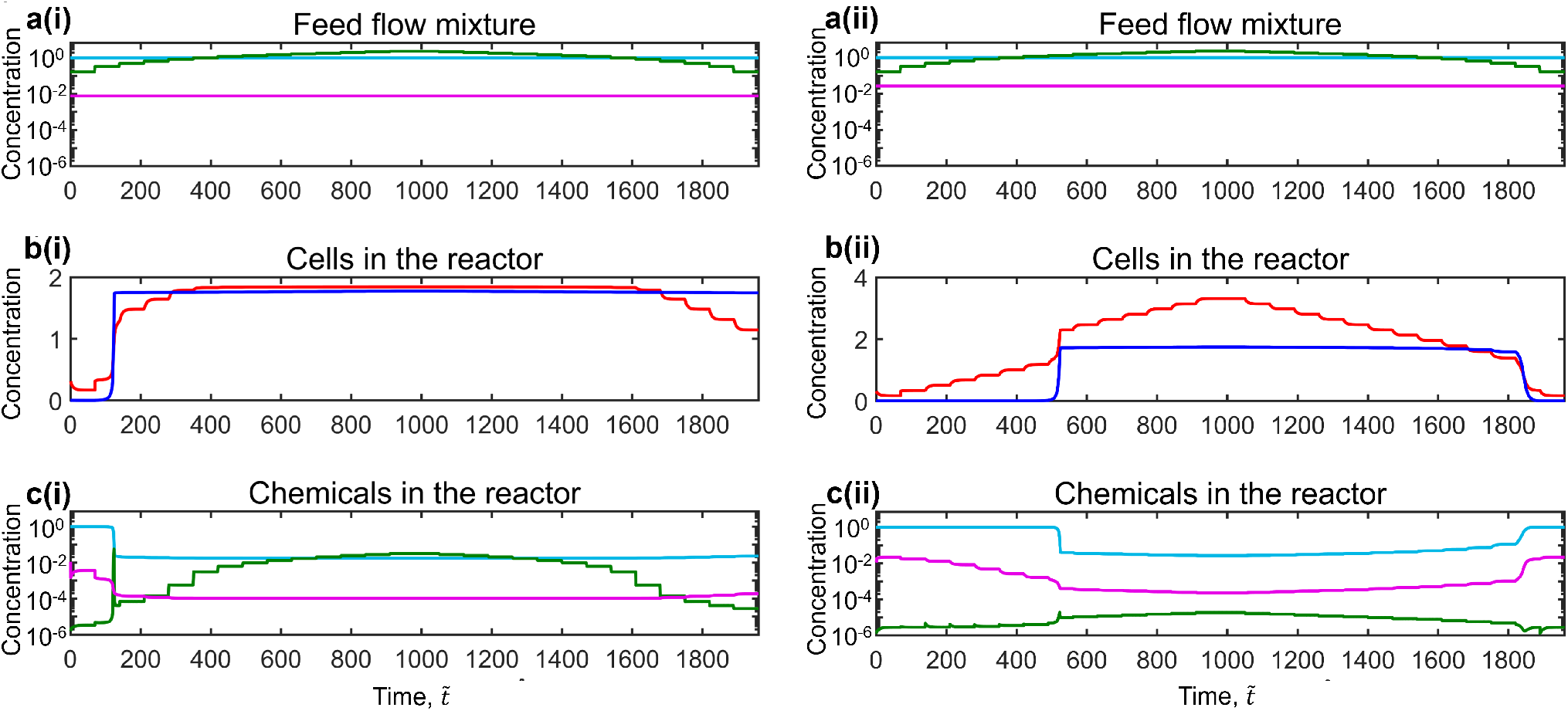
Removing the dependence of oxygen consumption by aerobes on carbon availability eliminates oscillatory growth dynamics and yields simulations that closely match prior metabolic model predictions. **(i)** Results of the identical simulation as in Fig. 1, but with *κ*_*o*,aer,min_ = *κ*_*o*,aer,max_ (*ϵ* = 1). In this case, because oxygen is depleted more rapidly, the coexistence state arises earlier and persists over the entire range of *β* that is subsequently explored. Because this range of *β* does not induce a transition from coexistence (C) back to the aerobes-only (AO) state, oscillations (which arise during this transition as shown in Fig. 1) do not have the chance to occur. To further probe this behavior, in **(ii)** we increase the value of oxygen inflow concentration to 43 μM (*δ* of 0.014), which induces the transition from the C to AO states. Nevertheless, in this case, oscillations still do not occur, indicating that the dependence of oxygen consumption on carbon availability is necessary for the oscillatory dynamics observed in Fig 1 to arise. Notably, **(ii)** indicates that the concentration of aerobes continually increases with *β* in the coexistence regime 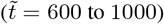 in this case, as was also observed in^2^; moreover, the regime over which bistability persists corresponds closely to the results of the more sophisticated metabolic modeling in^2^.

#### Bacterial growth

Simple sugar consumption then results in cellular growth. The rate at which anaerobe concentration increases has a contribution from the consumption of non-shared sugars, *N*(1 – Ξ)*b*_an_*γ*_an_*g*_*x*,an_(*x*)[1 – *g*_*o*,an_(*o*)], and another contribution from the consumption of liberated simple sugar that is shared among all bacteria, *b*_an_*γ*_an_*g*_*s*,an_(*s*)[1 – *g*_*o*,an_(*o*)]; here, *γ*_an_ is the maximal cellular growth rate per unit simple sugar. Conversely, the rate at which aerobe concentration increases only reflects the consumption of shared simple sugar, *b*_aer_*γ*_aer_*g*_*s*,aer_(*s*)*g*_*o*,aer_(*o*), where again *γ*_aer_ is the maximal cellular growth rate per unit simple sugar. This formulation of our model treats oxygen as a joint mediator of both aerobic and anaerobic metabolism of the sugar, albeit in opposing ways; an alternative formulation that instead employs Liebig’s law of the minimum, wherein the growth rate is set solely by the scarcest of the two (oxygen or sugar) yields identical results, as shown in Fig. 7.

**Figure 7.**
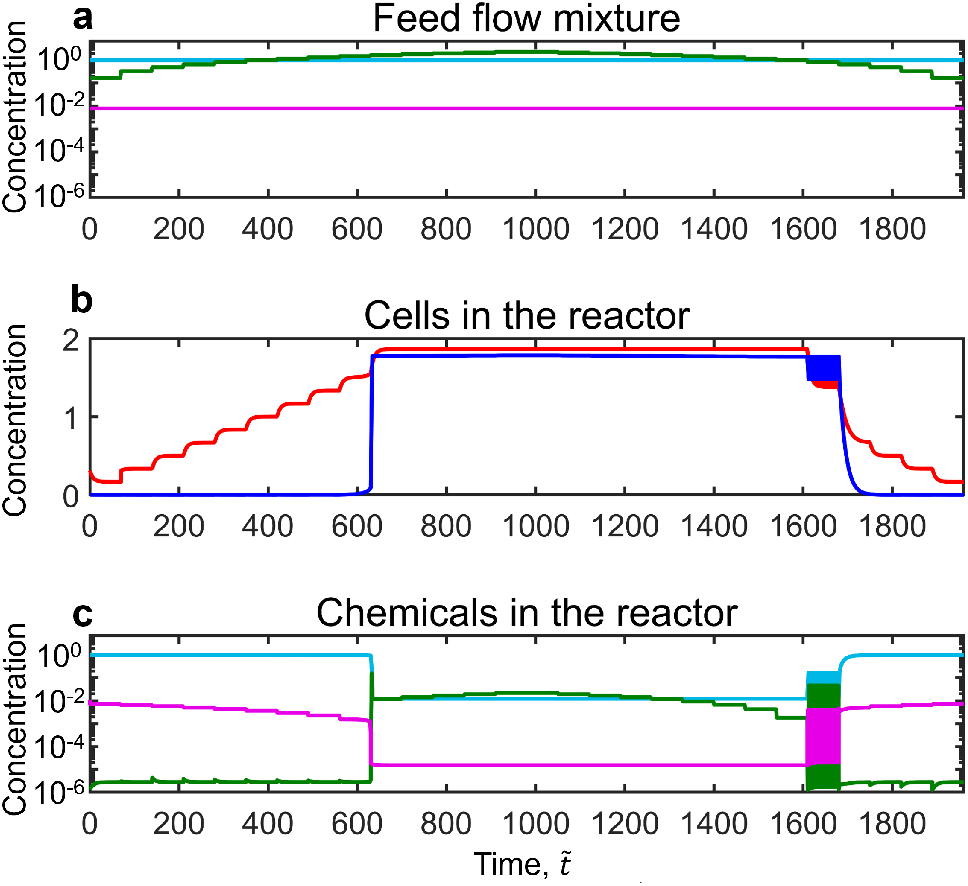
Our findings of bistability and oscillations are unchanged upon incorporating Liebig’s law of the minimum. For simplicity and to make our model more generally applicable to other mediators of microbial metabolism (e.g., pH, signaling molecules, toxins), we take oxygen as a joint mediator of carbon metabolism. Alternatively incorporating Liebig’s law of the minimum, wherein the scarcest of the two (oxygen or sugar) determines growth rate and the uptake rate of the non-limiting of the two, yields identical results as shown above (identical simulation as in Fig. 1 but with Liebig’s law). For this formulation, we define a function *g*_aer_(*o, s*) ≡ min{*g*_*s*,aer_(*s*), *ωg*_*o*,aer_(*o*)}, where *ω* = 6 is the utilization ratio that corresponds to the oxygen molecules required to fully utilize one glucose monomer^42^. Then, we replace *κ*_*o*,aer_(*s*)*g*_*o*,aer_(*o*) with *κ*_*o*,aer,max_*g*_aer_(*o, s*) in Eq. 3 and *g*_*s*,aer_(*s*)*g*_*o*,aer_(*o*) with *g*_aer_(*o, s*) in Eqs. 2 and 5.

#### Governing equations

Our model is thus summarized as:

Complex carbohydrate:

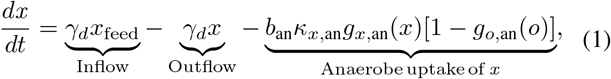

Simple sugar:

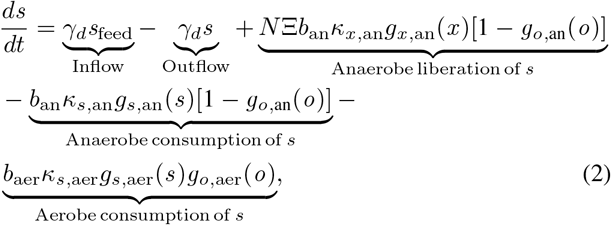

Oxygen:

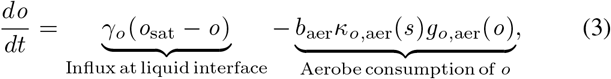

Anaerobes:

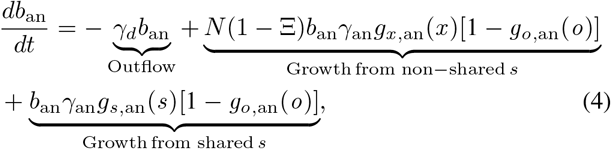

Aerobes:

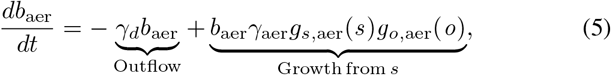

where *t* represents time.

### B. Characteristic dimensionless parameters

Non-dimensionalizing Eqs. 1–5 yields useful dimensionless parameters characterizing the biological, chemical, and physical dynamics underlying this complex system. In particular, we choose to rescale {*t, x, s, o, b*_an_, *b*_aer_} by the characteristic quantities 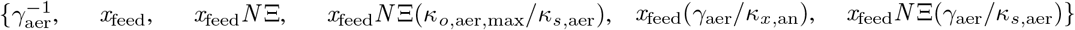 that describe the aerobe doubling time, feed concentration of complex carbohydrate, concentration of liberated simple sugar from this feed, corresponding concentration of consumed oxygen, corresponding concentration of newly-grown anaerobes, and corresponding concentration of newly-grown aerobes, respectively. We also rescale *κ*_*o*,aer_(*s*) by *κ*_*o*,aer,max_. This process yields the non-dimensional equations:

Complex carbohydrate:

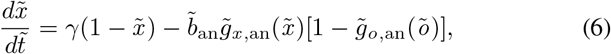

Simple sugar:

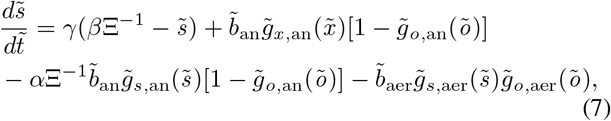

Oxygen:

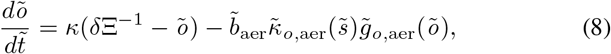

Anaerobes:

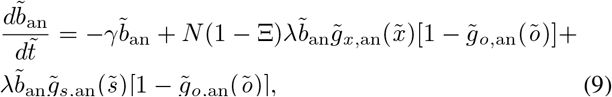

Aerobes:

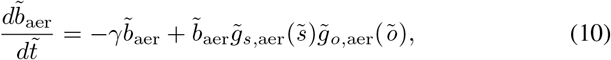

where the tildes indicate non-dimensionalized variables. Seven dimensionless parameters emerge:

- The *anaerobic uptake* parameter *α* ≡ *κ*_*s*,an_/(*κ*_*x*,an_*N*) compares the anaerobic carbon uptake rates of simple sugar and complex carbohydrates. When *α* > 1, anaerobe uptake of simple sugar is faster than that of complex carbohydrates, whereas when *α* < 1, anaerobe uptake of complex carbohydrates dominates. As shown in Table I, typical experiments can fall in either regime.
- The *simple sugar abundance* parameter *β* ≡ *s*_feed_/(*x*_feed_*N*) compares the abundance of simple sugar in the feed with that contained in the complex carbohydrate in the feed. When *β* > 1, carbon enters the system predominantly in the form of simple sugar, whereas when *β* < 1, carbon enters the system predominantly in the form of complex carbohydrates. As shown in Table I, typical experiments fall in this latter regime.
- The *aerobic washout* parameter *γ* ≡ *γ_d_*/*γ*_aer_ compares the rates of aerobic growth and dilution. When *γ* > 1, aerobes are diluted from the reactor faster than they can grow, leading to their washout, whereas when *γ* < 1, aerobic growth is sufficiently fast for them to remain in the reactor. As shown in Table I, typical experiments fall in this latter regime.
- The *oxygen abundance* parameter *δ* ≡ (*o*_sat_/*κ*_*o*,aer,max_)/ (*x*_feed_*N*/*κ*_*s*,aer_) compares the time needed for aerobes to deplete oxygen from the saturated influx with the time needed for them to deplete the quantity of simple sugar contained in the feed of complex carbohydrate. When *δ* > 1, depletion of oxygen by the aerobes is slow compared to that of simple sugar, whereas when *δ* < 1, oxygen is rapidly depleted by aerobic consumption. As shown in Table I, typical experiments fall in this latter regime.
- The *oxygen transfer* parameter *κ* ≡ *γ_o_*/*γ*_aer_ compares the rates of oxygen influx and aerobic growth. When *κ* > 1, the oxygen supply is rapidly replenished to its saturation concentration *o*_sat_, whereas when *κ* < 1, oxygen influx is slow. As shown in Table I, typical experiments fall in the former regime.
- The *growth* parameter *λ* ≡ *γ*_an_/*γ*_aer_ compares the rates of anaerobic and aerobic growth. When *λ* > 1, anaerobes can grow much faster than aerobes, whereas when *λ* < 1, aerobic growth dominates. As shown in Table I, typical experiments can fall in either regime.
- The *oxygen consumption* parameter *ϵ* ≡ *κ*_*o*,aer,min_/*κ*_*o*,aer,max_ compares the minimum and maximum oxygen consumption rates by aerobes under nutrient-depleted or nutrient-replete conditions, respectively. When *ϵ* =1 (its maximal value, by definition), oxygen consumption by aerobes does not depend on simple sugar availability, whereas when *ϵ* ≪ 1 oxygen consumption is strongly dependent on simple sugar levels. As shown in Table 1, in this work, we consider both limits for generality.

**Table I.** Dimensionless parameters characterizing our model. Also listed are ranges of their values reported in previous studies.

| Parameter | Name | Values | References |
| --- | --- | --- | --- |
| $\Xi$ | Shared fraction | 0 to 1 | 2,5,13,93,94 |
| $N$ | Degree of polymerization | $10^0$ to $10^2$ | 2,5,13,93,94 |
| $\alpha \equiv \kappa_{s,\text{an}}/(\kappa_{x,\text{an}}N)$ | Anaerobic uptake | 0 to $10^1$ | 2,13,89,93 |
| $\beta \equiv s_{\text{feed}}/(x_{\text{feed}}N)$ | Simple sugar abundance | 0 to $10^0$ | 2,3,5,7,13,89,93,103,104 |
| $\gamma \equiv \gamma_d/\gamma_{\text{aer}}$ | Aerobic washout | 0 to $10^0$ | 2,3,5,89,93,103–105 |
| $\delta \equiv (o_{\text{sat}}/\kappa_{o,\text{aer},\text{max}})/(x_{\text{feed}}N/\kappa_{s,\text{aer}})$ | Oxygen abundance | $10^{-3}$ to $10^{-1}$ | 2,42,89,102 |
| $\kappa \equiv \gamma_o/\gamma_{\text{aer}}$ | Oxygen transfer | $10^1$ to $10^2$ | 2,105 |
| $\lambda \equiv \gamma_{\text{an}}/\gamma_{\text{aer}}$ | Growth | 0 to $10^1$ | 5,13,55,94,103,104 |
| $\epsilon \equiv \kappa_{o,\text{aer},\text{min}}/\kappa_{o,\text{aer},\text{max}}$ | Oxygen consumption | 0 to $10^0$ | See Fig. 6 |

Our overarching goal is to address the question: *For a given microbial community, how does changing the balance of carbon and oxygen fluxes alter its overall composition?* To this end, in what follows, we use the model summarized by Eqs. 6–10 to explore the influence of varying *β* and *δ*, keeping all other parameters constant.

### C. Implementation of numerical simulations

First, to explore the influence of varying simple sugar abundance, we systematically vary sfeed (which varies *β*) and use numerical simulations of Eqs. 6–10 to examine the resulting variations in the complex carbohydrate, simple sugar, oxygen, aerobe, and anaerobe concentrations in the reactor, representedby the non-dimensional variables 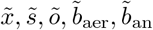, respectively. We keep all the other input parameters constant at their representative experimental values as given in Table II. Then, to explore the additional influence of varying oxygen depletion, we also vary *o*_sat_ (which varies *δ*) and examine the resulting variations in chemical and bacterial concentrations. Finally, to investigate the pivotal influence of metabolic mutualism, we examine these behaviors for different values of the sharing parameter Ξ, ranging from a high value of 0.9 representing “altruistic” liberation of sugars by the anaerobes to a low value of 0.1 representing “greedy” utilization of sugars by the anaerobes.

**Table II.** Parameter values used in our simulations. Upper and lower tables show dimensional and corresponding dimensionless parameters, respectively. Values not listed are the same as in Fig. 1. * This value was chosen to be much lower than the aerobic growth rate, while still being non-zero.

| Parameter | Value in Fig. 1 | Value in Fig. 2 | Value in Fig. 3 | References |
| --- | --- | --- | --- | --- |
| $\Xi$ | 0.5 | | 0.1, 0.5, 0.9 | <sup>2</sup> |
| $N$ | 100 | | | <sup>2</sup> |
| $\Delta t$ | 50 h | 200 h | 200 h | <sup>2</sup> |
| $o_{\text{sat}}$ | 13 $\mu\text{M}$ | | 1.8 to 23 $\mu\text{M}$ | <sup>2</sup> |
| $x_{\text{feed}}$ | 60 $\mu\text{M}$ | | | <sup>2</sup> |
| $s_{\text{feed}}$ | 0.5 to 7 mM | | | <sup>2</sup> |
| $b_{\text{aer},0}$ | $1.7 \times 10^8$ cells/mL | | | <sup>2</sup> |
| $b_{\text{an},0}$ | $5.0 \times 10^6$ cells/mL | | | <sup>2</sup> |
| Initial $s$ | 0 | | | <sup>2</sup> |
| Initial $o$ | 18 $\mu\text{M}$ | | | |
| Initial $x$ | 60 $\mu\text{M}$ | | | <sup>2</sup> |
| $o_{1/2,\text{aer}}$ | 1 $\mu\text{M}$ | | | <sup>105</sup> |
| $o_{1/2,\text{an}}$ | 0.35 $\mu\text{M}$ | | | <sup>2</sup> |
| $x_{1/2,\text{an}}$ | 8 $\mu\text{M}$ | | | <sup>2</sup> |
| $s_{1/2,\text{aer}}$ | 0.05 $\mu\text{M}$ | | | <sup>2</sup> |
| $s_{1/2,\text{an}}$ | 1.4 mM | | | <sup>2</sup> |
| $\kappa_{o,\text{aer},\text{max}}$ | $1.2 \times 10^{-12}$ mM (cells/mL)<sup>-1</sup> s<sup>-1</sup> | | | <sup>105</sup> |
| $\kappa_{o,\text{aer},\text{min}}$ | $3.3 \times 10^{-14}$ mM (cells/mL)<sup>-1</sup> s<sup>-1</sup> | | | <sup>42,102</sup> |
| $\kappa_{x,\text{an}}$ | $6.7 \times 10^{-15}$ mM (cells/mL)<sup>-1</sup> s<sup>-1</sup> | | | <sup>2</sup> |
| $\kappa_{s,\text{aer}}$ | $2.2 \times 10^{-12}$ mM (cells/mL)<sup>-1</sup> s<sup>-1</sup> | | | <sup>2</sup> |
| $\kappa_{s,\text{an}}$ | $9.1 \times 10^{-13}$ mM (cells/mL)<sup>-1</sup> s<sup>-1</sup> | | | <sup>2</sup> |
| $\gamma_{\text{aer}}$ | 1.4 h<sup>-1</sup> | | | <sup>42</sup> |
| $\gamma_{\text{an}}$ | 0.05 h<sup>-1</sup> | | | * see caption |
| $\gamma_o$ | 47.4 h<sup>-1</sup> | | | <sup>2</sup> |
| $\gamma_d$ | 0.2 h<sup>-1</sup> | | | <sup>2</sup> |
| $\Xi$ | 0.5 | | 0.1, 0.5, and 0.9 | |
| $N$ | 100 | | | |
| $\alpha$ | 1.4 | | | |
| $\beta$ | 0.083 to 1.2 | | | |
| $\gamma$ | 0.14 | | | |
| $\delta$ | 0.0039 | | 0.00056 to 0.0072 | |
| $\kappa$ | 34 | | | |
| $\lambda$ | 0.036 | | | |
| $\epsilon$ | 0.029 | | | |
| $\tilde{o}_{1/2,\text{aer}}$ | $6.2 \times 10^{-4}$ | | | |
| $\tilde{o}_{1/2,\text{an}}$ | $2.2 \times 10^{-4}$ | | | |
| $\tilde{x}_{1/2,\text{an}}$ | 0.13 | | | |
| $\tilde{s}_{1/2,\text{aer}}$ | $1.7 \times 10^{-5}$ | | | |
| $\tilde{s}_{1/2,\text{an}}$ | 0.47 | | | |

To do so, we use the `ode23t` solver in `MATLAB`, initializing each simulation with low aerobe and anaerobe concentrations *b*_aer,0_ and *b*_an,0_ chosen to be ≈ 0.3 and ≈ 10^−3^ times smaller than the characteristic concentrations {*x*_feed_(*γ*_aer_/*κ*_*x*,an_), *x*_feed_*N*Ξ(*γ*_aer_/*κ*_*s*,aer_)} (Table II), matching the experiments in^2^. The initial time step is 10^−15^ s, chosen to be > 10^18^ times smaller than the characteristic time scale 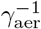. The solver then varies the time step dynamically to ensure a relative error tolerance of 10^−4^ for all quantities or absolute error tolerances of 10^7^ cells/mL and 10^−6^ μM for cells and chemicals, respectively; these tolerance values are chosen to be ≈ {50, 400} and ≈ {60, 3000, 1600} times smaller than the characteristic concentrations {*x*_feed_(*γ*_aer_/*κ*_*x*,an_), *x*_feed_*N*Ξ(*γ*_aer_/*κ*_*s*,aer_)} (cells) and {*x*_feed_, *x*_feed_*N*Ξ, *x*_feed_*N*Ξ(*κ*_*o*,aer,max_/*κ*_*s*,aer_)} (chemicals), respectively. For each distinct {*β, δ*} condition explored, we then solve Eqs. 6–10 for a duration of 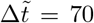 to match2 (Fig. 1) or 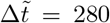 (Figs. 2–3) to ensure we reach a (possibly dynamic) steady state, using the final state of the preceding condition to initialize the next with a new value of *β*, representing a change in the influx of fresh simple sugar into the reactor. Furthermore, as in^2^, we also employ a “reinoculation protocol” in which if either {*b*_aer_, *b*_an_} drops below the low initial concentrations {*b*_aer,0_, *b*_an,0_} at the end of a given simulation condition, we reset them to those initial values at the beginning of the next simulation condition tested to prevent irreversible washout of cells from the reactor.

## III. RESULTS

### A. Chemical and bacterial dynamics under varying simple sugar abundance

How do variations in simple sugar abundance alter the composition of the overall microbial community? And are these alterations history dependent, as is often observed in experiments? To answer these questions, we first investigate the coupled chemical and bacterial dynamics quantified by Eqs. 6–10 that emerge upon successive variations in the feed simple sugar concentration *s*_feed_ (i.e., varying *β*) for an exemplary case of Ξ = 0.5 and *N* = 100. In particular, as shown in Fig. 1b, we hold the feed concentrations of oxygen (magenta) and complex carbohydrate (teal) constant, and successively increase sfeed (green, increasing *β*) step-wise every 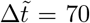 (“ramp up”). To investigate any possible history dependence, we then “ramp down” by successively decreasing sfeed (decreasing *β*) step-wise by the same amount. The resulting concentrations of cells and chemicals in the reactor are shown in Figs. 1c–d, respectively.

#### The aerobes-only state

During the initial phase of the ramp up period (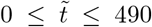, denoted by I in Fig. 1c), oxygen consumption by the aerobes is insufficient to sustain anaerobic growth (dark blue, Fig. 1c). As a result, the microbial community is in the *aerobes-only* (AO) state in which solely aerobes consume the supplied simple sugar (green, Fig. 1d) and utilize it for their growth; the concentration of complex carbohydrate (teal) remains unchanged, since this is only taken up by anaerobes. In this state, with each successive increase in simple sugar abundance *β*, the aerobe concentration exponentially reaches a new steady-state value (red, Fig. 1c). The oxygen concentration concomitantly approaches progressively decreasing steady-state values due to aerobic consumption (magenta, Fig. 1d). For clarity of notation, we denote long-time steady-state values of chemical and bacterial concentrations by the subscript s hereafter.

#### Coexistence and hysteresis

As the concentration of aerobes rises with increasing simple sugar abundance *β*, the oxygen concentration in the reactor eventually decreases enough to sustain anaerobic growth as well—as shown by the drop in the magenta line just before 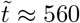 in Fig. 1d. Consequently, the anaerobe concentration increases dramatically (dark blue, Fig. 1c); we denote this transition by II.

As the anaerobes proliferate, they take up the exogenously-supplied complex carbohydrate (teal, Fig. 1d) and continually liberate new simple sugar to be shared by the entire microbial community (Fig. 8). Despite the low oxygen concentration in the reactor, aerobes can also proliferate, albeit with low metabolic activity. Hence, these carbon-replete conditions give rise to the *coexistence* (C) state (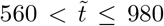, denoted by III in Fig. 1c). Both aerobes (red) and anaerobes (dark blue) are maintained at nearly constant concentrations, buffered against subsequent changes in the feed simple sugar supply.

**Figure 8.**
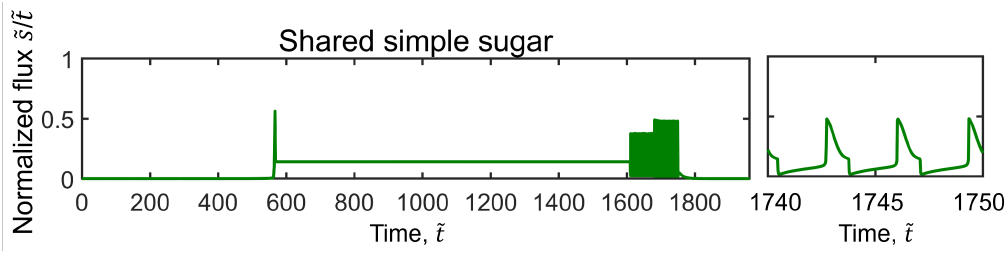
Dynamics of simple sugar sharing by the anaerobes. The shared flux of simple sugar is given by 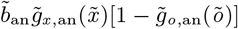 (the second term in Eq. 7) and is shown for the simulation in Fig. 1. On the right, we show a magnified view of the oscillatory dynamics immediately before the transition from the C to AO states.

Indeed, the coexistence state also persists as the simple sugar abundance *β* is ramped down (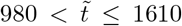, denoted by IV in Fig. 1c). Strikingly, we observe hysteretic behavior: the chemical and bacterial concentrations during the ramp down do not mirror those of the ramp up period—as observed experimentally as well^2^. For example, the coexistence state persists over a broader range of simple sugar abundance *β*, including at values that were too low to initiate coexistence during ramp up; see, for example, 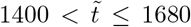 in Fig. 1c–d. Having established a mutually-beneficial state, this mixed microbial community continues to coexist, despite the decreasing levels of exogenously-supplied simple sugar.

#### Oscillatory dynamics

Further decreasing the simple sugar abundance *β* leads to a new mode of coexistence. Surprisingly, we observe sustained oscillations in both chemical and bacterial concentrations (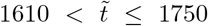, denoted by V in Fig. 1c)—again recapitulating some experimental observations^27^. The concentrations of both aerobes and anaerobes rise and fall in phase (magnified view to the right of Fig. 1c). The chemical concentrations correspondingly switch between metastable conditions resembling the coexistence state with low oxygen, but high simple sugar availability, and the aerobes-only state with high oxygen, but low simple sugar availability (Fig. 1d, right).

These complex dynamics again reflect the central role of microbial mutualism. In the low-oxygen metastable state, anaerobes can take up complex carbohydrate (teal) and liberate new simple sugar molecules that are utilized by the entire community (e.g., rise in the green line at 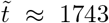). Then, as the complex carbohydrate is increasingly depleted by the anaerobes, less simple sugar is liberated (Fig. 8) and available for the community to use (drop in the green line for 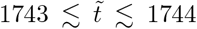). Because oxygen consumption depends on carbon availability, aerobic consumption of oxygen becomes concomitantly limited—eventually driving an abrupt transition to the high-oxygen metastable state (e.g., rise in the magenta line at 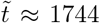). Under these conditions, anaerobic metabolism is impeded but not completely abrogated, enabling the concentration of complex carbohydrate to gradually be replenished (teal, 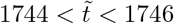) and broken down into shared simple sugar. Thus, the amount of shared simple sugar concomitantly increases (Fig. 8), enabling the aerobes to increasingly deplete oxygen (magenta, 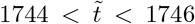)—eventually driving another abrupt transition back to the low-oxygen state 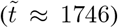 in which anaerobic growth is sustained again. Indeed, removing the dependence of oxygen consumption by aerobes on simple sugar availability abolishes the oscillations entirely (Fig. 6)—providing further confirmation of this picture.

#### Transition back to the aerobes-only state

Further decreasing the simple sugar abundance *β* eventually causes the high-oxygen state to dominate—leading to a precipitous decline in the concentration of anaerobes (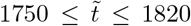, denoted by VI in Fig. 1c). The community thus transitions back to the AO state denoted by VII in Fig. 1c.

Taken together, these results (Fig. 1b–d) highlight the rich chemical and bacterial behaviors that emerge from our minimal model of metabolic mutualism. Remarkably, despite the simplicity of the model compared to those that consider a more complex network of metabolic fluxes, it successfully recapitulates the two different states observed in experiments upon varying carbon and oxygen fluxes^2^: the aerobes-only (AO) state and the state of aerobe-anaerobe coexistence (C). Our model also recapitulates the hysteretic nature of transitioning between these two states^2^ as well as the possible emergence of oscillations between them^27^, as observed in many experiments. Having recapitulated these behaviors, we next set out to uncover the underlying biophysical principles that govern them.

### B. Biophysical principles governing the different states and transitions between them

To summarize the findings in Fig. 1, we plot the steadystate bacterial concentrations *b*_aer,s_ and *b*_an,s_ for the different values of simple sugar abundance *β* tested in Fig. 2a–b; the corresponding steady-state concentrations of oxygen, complex carbohydrate, and simple sugar are shown in c–e. The circles show the results of the full numerical simulations, while the curves show the results of the theoretical predictions developed hereafter. The filled circles show the states that arise during the ramp up period (increasing *β*), while the open circles show the ramp down period (decreasing *β*). As described in §IIIA, the microbial community is initially in the aerobes-only state with *b*_aer,s_ > 0 and *b*_an,s_ = 0 (leftmost points in Fig. 2). As the simple sugar abundance *β* is increased, the aerobe concentration *b*_aer,s_ monotonically increases (denoted by I in panel a) while the anaerobe concentration *b*_an,s_ remains negligible. Then, above a critical value of *β* = *β*_aer,crit_ ≈ 0.6, the community abruptly transitions to the state of coexistence (II) with *b*_aer,s_ ≈ 1.85 and *b*_an,s_ ≈ 1.75, independent of subsequent increases in *β* (III). The reverse behavior is hysteretic upon ramping down: the community persists in the coexistent state over a broader range of 1.2 > *β* > 0.4 (IV), eventually exhibiting oscillatory dynamics (indicated by V and the undulating red line in panel a). It then abruptly transitions back to the aerobes-only state as simple sugar abundance decreases below a critical value *β* = *β*_coex,min_ ≈ 0.28 (VI–VII).

Having characterized these states and transitions between them, we next seek biophysical principles that quantitatively determine the:

- Steady-state concentration *b*_aer,s_ in the AO state (I and VII),
- Critical value *β*_aer,crit_ that determines the transition from the AO to C state (II),
- Steady-state concentrations *b*_aer,s_ and *b*_an,s_ in the C state (III and IV),
- Critical value *β*_coex,min_ that determines the subsequent transition from the C to AO state (VI).

#### The aerobes-only state

In the AO state, at each imposed value of the simple sugar abundance *β*, the aerobe concentration reaches a well-defined steady-state value *b*_aer,s_ (I and VII in Figs. 1c, 2); the simple sugar and oxygen concentrations reach corresponding steady-state values 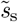 and 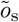, respectively. Therefore, we begin by seeking steady-state solutions to Eqs. 6–10 by setting 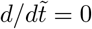. This procedure yields

From Eq. 6:

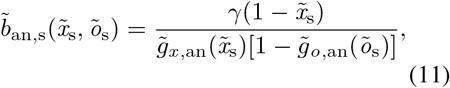

From Eq. 7:

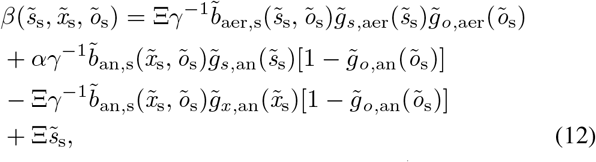

From Eq. 8:

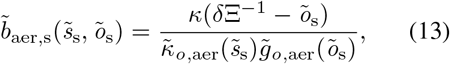

From Eq. 10:

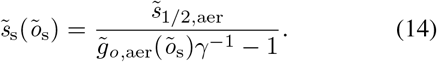

In the AO state, 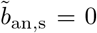; thus, Eq. 11 yields 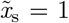. Substituting this equality combined with Eq. 14 into Eqs. 12-13 then yields both *β* and 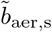 as functions of solely 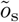. Inverting 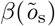 then yields our final result, the aerobes-only solutions for 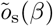 and 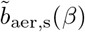 shown by the red curves in Fig. 2a,c. These solutions capture our numerical data (circles for I and VII) exactly—quantifying the intuition that in the AO state, aerobes consume the supplied simple sugar, mediated by oxygen consumption, and utilize it for their growth.

#### Transition to coexistence

As the oxygen concentration in the reactor decreases with increasing aerobe concentration, it eventually becomes low enough to sustain anaerobic growth as well (II in Figs. 1c–d, 2). Consequently, the microbial community transitions to the C state. Intuitively, this transition is analogous to the process by which a small inoculum of a new species (anaerobes) “invades” an existing colony of an indigenous species (aerobes)^54^—the onset of which is determined by the transition between 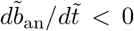 (anaerobic washout) and 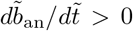 (anaerobic invasion). Therefore, we begin by setting 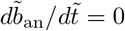, which yields

From Eq. 9:

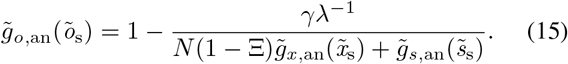

In the AO state immediately prior to the transition to coexistence, 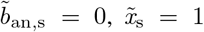, and 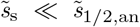 (Fig. 1d), and therefore, 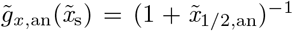 and 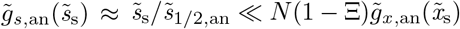. Substituting these simplifications into Eq. 15 then yields estimates for the critical oxygen concentration and corresponding simple sugar abundance at which the microbial community transitions to the C state: 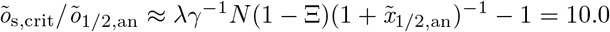 and 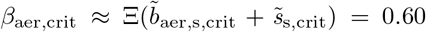, respectively, where 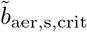 and 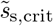 are given by substituting 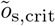 into Eqs. 13–14. These estimates are in excellent agreement with the values 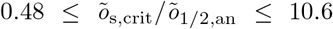 and 0.58 ≤ *β*_aer,crit_ ≤ 0.75 obtained from our numerical simulation (Figs. 1d, 2)—quantifying the intuition that to transition to the C state, oxygen consumption by aerobes must be sufficient to provide conditions for anaerobes to begin to “invade” the reactor without being washed out. For 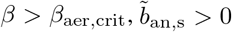, and therefore, the AO solution indicated by the red lines in Fig. 2 becomes unstable (dashed).

#### The coexistence state

Without any simplifications, the steady-state relations obtained previously (Eqs. 11–15) jointly yield exact analytical solutions for 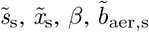, and 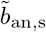 as functions of solely 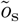. Therefore, we expect that these equations fully describe the steady-state chemical and bacterial concentrations in the C state. To verify this expectation, we plot these quantities for varying 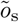, as shown by the blue curves in Fig. 2. The solutions capture our numerical data (circles for III and IV) exactly—quantifying the intuition that in the C state, anaerobes break down complex carbohydrate to simple sugar, sustaining their own growth as well as that of the aerobes, which also consume oxygen and thereby maintain favorable conditions for the anaerobes to continue to survive.

#### Transition back to the aerobes-only state

Upon ramping down the simple sugar abundance *β*, coexistence persists until *β* = *β*_coex,min_ < *β*_aer,crit_: the extant population of anaerobes shares simple sugar with the aerobes, enabling them to continue to grow, consume oxygen, and sustain anaerobic growth as well—despite the low value of *β*. The critical value *β*_coex,min_ is then simply given by the minimum value of *β* for which a real-valued solution to the coexistence state (as obtained earlier, shown by the blue curves in Fig. 2) exists. We thereby estimate *β*_coex,min_ ≈ 0.29, in excellent agreement with the value 0.25 ≤ *β*_coex,min_ ≤ 0.33 obtained from our numerical simulation—quantifying the intuition that even lower levels of simple sugar abundance do not enable aerobes to consume enough oxygen to sustain anaerobic growth. As a result, the C solution indicated by the blue lines in Fig. 2 becomes unstable (dashed) and the community transitions back to the AO state (VI). Within the context of dynamical systems theory, the oscillatory dynamics that arise prior to this transition (V) are characteristic of a limit cycle around a stable equilibrium point in phase space; indeed, the amplitude of the oscillations in e.g., oxygen concentration displays a hysteresis loop characteristic of a subcritical Hopf bifurcation, as shown in the inset to Fig. 2c. Further investigating the features of this instability will be an interesting direction for future work.

Taken together, these results demonstrate that not only does our full model recapitulate the two states (AO and C) observed in experiments, as well as the hysteretic transitions between them, but the steady-state solutions given by Eqs. 11–15 also provide a way to analytically describe these behaviors. Thus, to develop a broader understanding of the underlying biophysics, we next examine how these complex phenomena depend on both oxygen depletion and sharing of simple sugar more generally.

### C. Oxygen depletion and nutrient sharing jointly enable coexistence

Given the necessity of oxygen depletion for coexistence, we hypothesize that increasing *δ*—which corresponds to reduced depletion of oxygen by the aerobes—will shift the AO-C transition to larger values of simple sugar abundance *β*. To test this hypothesis, we repeat the simulations shown in Fig. 2, ramping *β* up and then down between 0 and 1.2, keeping the same intermediate value of the sharing parameter Ξ = 0.5 but with varying *δ*. The results are shown in the state diagram shown in the middle panel of Fig. 3, with the results of Figs. 1–2 given by the points along *δ* = 3.9 × 10^−3^. Consistent with the findings reported in Figs. 1–2, the microbial community initially exists in the aerobes-only (AO) state (red triangles) at low *β*, eventually transitioning to the coexistence (C) state (blue circles) as *β* is ramped up above a critical value *β*_aer,crit_. Moreover, as expected for the case of large *δ*—for which *β*_aer,crit_ exceeds the upper limit of the *β* explored—coexistence is never reached and the bistable regime is precluded. Upon ramping down *β*, the community persists in the C state in a bistable regime (superimposed red triangles and blue circles) until it eventually transitions back to the AO state below another critical value *β*_coex_,min. In agreement with our hypothesis, these states and the transitions between them are sensitive to oxygen depletion: as oxygen depletion by the aerobes is reduced (increasing *δ*), the AO state persists over a broader range of *β*, with both transitions between the AO and C states shifted to larger values of *β*. Our analytical steady-state solutions given by Eqs. 11–15 capture these shifts exactly; the shading in Fig. 3 corresponds to the top bar in Fig. 2, indicating the AO and C states as well as the bistable regime between them, and the dashed lines show the analytical solutions for *β*_aer,crit_ (red) and *β*_coex,min_ (blue). This close agreement quantifies the intuition that coexistence requires both oxygen depletion and nutrient sharing.

As a final demonstration of this point, we repeat these simulations, but for different values of the sharing parameter Ξ, as well. In particular, we explore two different limits: that of large Ξ = 0.9, in which the anaerobes are “altruistic” and share a larger fraction of the simple sugar liberated from complex carbohydrate with the entire community, and conversely that of small Ξ = 0.1, in which the anaerobes are instead “greedy” and only share a smaller fraction of the liberated simple sugar. The results are summarized by the left- and right-most panels of Fig. 3, respectively; the theoretical predictions for the resulting states and transitions between them, as determined using Eqs. 11–15, are shown by the shaded regions and dashed lines demarcating them, respectively. We find excellent agreement between the full numerical simulations and the analytical theory, confirming the validity of our biophysical analysis more broadly. In both cases of altruistic or greedy anaerobes, we again find that the overall community initially exists in the AO state at low simple sugar abundance *β*, eventually transitions to the C state with increasing *β*, and then continues to coexist with decreasing *β* until it transitions back to the AO state at even lower *β*. Moreover, we again find that the AO (C) state spans a wider range of *β* at larger (smaller) *δ*, highlighting the importance of oxygen depletion in enabling coexistence.

Confirming our expectation, nutrient sharing also plays a pivotal role in enabling coexistence. For example, in the altruistic case of Ξ = 0.9, coexistence persists over a broader window of {β, *δ*} after having been established i.e., during ramp down, as shown by the left-most panel of Fig. 3. Conversely, in the greedy case of Ξ = 0.1, the regime of bistability over which coexistence can persist nearly vanishes, as shown by the right-most panel. Close examination of the corresponding shifts in the boundaries between the AO and C states *β*_aer,crit_ (red) and *β*_coex,min_ (blue) reveals the underlying cause. Increased sharing of simple sugar (larger Ξ) increases *β*_aer,crit_ (red line shifts to the right): it is more difficult to initially sustain the population of anaerobes if they utilize less of the complex carbohydrate for their own growth. However, despite this cost associated with altruism, it can also be beneficial. Increased sharing also decreases *β*_coex,min_ (blue line shifts to the left): having been stably established in the community, anaerobes can continue to supply simple sugar to sustain aerobic growth, which in turn maintains the low-oxygen environment needed for the anaerobes themselves to continue to survive. Thus, as the old adage goes, *sharing is caring*.

## IV. DISCUSSION AND CONCLUSION

Centuries of research have established that competition for limited resources—the “frequently recurring struggle for existence”, as Charles Darwin put it^106^—can profoundly impact the growth and functioning of a multi-species community. It is now becoming increasingly clear that mutualistic interactions between distinct cell types also abound and play key roles in microbial communities; however, the dizzyingly-complex array of different interactions that arise in nature makes it challenging to isolate the influence of mutualism on community behavior. To address this challenge, in this work, we examined the growth of a simplified aerobe-anaerobe community with mutualistic metabolic interactions between the two species: the anaerobes sustain aerobic growth by breaking down non-metabolized complex carbohydrate to shared simple sugar, while the aerobes sustain anaerobic growth in turn by consuming oxygen.

Remarkably, despite its simplicity, we found that our model of this community recapitulates many of the behaviors exhibited by laboratory and natural communities. In particular, it can adopt *multiple stable states*—transitioning between the states of aerobe-only (AO) growth and aerobe-anaerobe coexistence (C) in response to changes in carbon and oxygen fluxes, as summarized in Fig. 4. It also exhibits *multistability*—either of the AO and C states can arise under identical conditions, depending on the history of carbon and oxygen fluxes—leading to *hysteresis* and even *oscillatory* growth dynamics. Taken together, these results provide a simple demonstration of how mutualism can generate the complex growth behaviors that commonly arise in many different microbial communities. As such, our work complements the vast and insightful literature in microbial ecology focusing on the role of competition in influencing community behavior. Examining the influence of additional non-mutualistic interactions in our model would therefore be an interesting extension of work.

Indeed, a strength of our model is that it can reproduce the community behavior predicted by a more sophisticated metabolic model of a similar aerobe-anaerobe community^2^ (as detailed further in Fig. 6). This more sophisticated model considers the full genome-scale network—comprising thousands of different metabolites and reactions—of metabolic interactions, which span the range from competitive to mutualistic. By contrast, ours only considers the interactions shown in Fig. 1a(ii), which are primarily mutualistic for the parameter values we used. This agreement between the model predictions could simply be fortuitous; but if not, it would corroborate the idea that consumption and secretion of only a small number of metabolites often dominates community behavior, as suggested by some experiments^3–5,7,13,15,94^. Further study is needed to test this idea and disentangle the relative contributions of different metabolites/metabolic pathways on community-level functioning.

Another strength of our model is that, owing to its simplicity, it provides analytical predictions (Eqs. 11–15) and biophysical intuition for the different behaviors exhibited by this microbial community. Our approach could thus be used to inform the design of future experiments seeking to better characterize these behaviors, without requiring the large quantities of data—which may not be readily accessible—that are needed as inputs to sophisticated metabolic models. For example, the analytical predictions accurately describe the chemical and bacterial concentrations in both the AO and C states, as well as the transitions between the states (Figs. 2–3), governed by the interplay of both oxygen depletion and nutrient sharing. In particular, our analysis shows that for aerobe-anaerobe coexistence to arise, aerobes must be able to consume sufficient oxygen to provide conditions for anaerobes to begin to “invade” the reactor. It also reveals the origins of hysteresis when transitioning away from coexistence: anaerobe sharing of simple sugar with the aerobes enables them to continue to grow and consume oxygen, even in conditions of reduced simple sugar supply, thereby sustaining coexistence. Finally, our analysis demonstrates that oscillations in bacterial growth—which are usually thought to be caused by competition^8,13,19,27–33^—arise in this community instead due to periodic fluctuations in oxygen depletion by the aerobes that are coupled to fluctuations in carbon availability, mediated by anaerobic growth and metabolism. Consistent with this interpretation, removing the dependence of oxygen consumption by the aerobes on carbon availability—as may sometimes be the case^2^—abolishes the oscillations (Fig. 6). Guided by our findings, it would be interesting to further explore the nature of these oscillations in experiments, in addition to further characterizing them within the framework of dynamical systems theory.

The community considered here provides a simplified representation of the aerobe-anaerobe communities that play critical roles in human health, our environment, waste treatment, and other biotechnological processes^2,4,5,7,11,39–55,65–74,76–88^; thus, our work sets a foundation for future research with potential implications extending beyond microbiology and biological physics. To this end, our model takes a step toward capturing the essential biophysical processes underlying the complex dynamics of such microbial communities—but in doing so, necessarily required some simplifying assumptions and approximations. For example, we considered growth of two species in the well-mixed environment of a bioreactor, in which the distribution of all chemicals is spatially uniform, and the imposed oxygen and carbon fluxes are held constant until the community reaches a steady state. However, natural environments often have spatial and temporal fluctuations in nutrient availability^107,108^, involving a larger repertoire of metabolites and mediators of metabolism, and with more than two microbial species that may adapt to changing conditions through evolutionary adaptation or phenotypic plasticity. These added complexities can give rise to fascinating new behaviors, leading to e.g., the emergence of spatial structure and facilitated coexistence in communities^3,4,19,70,89,109,110^. It could also be that microbes tune the extent of sharing (quantified by Ξ), which presumably could be under evolutionary selection, to e.g., optimize their individual growth or promote coexistence. Investigating these effects using the foundation provided by our model will be an important direction for future research.

## V. ACKNOWLEDGEMENTS

It is a pleasure to acknowledge Ned Wingreen for stimulating discussions and helpful feedback on the manuscript, as well as R. Kōnane Bay, Sebastian Gonzalez La Corte, Anna Hancock, Carolina Trenado Yuste, and Hongbo Zhao for useful conversations. This work was supported by the Princeton Catalysis Initiative, the Pew Biomedical Scholars Program, NSF grant EF-2124863, and in part by funding from the Princeton Center for Complex Materials, a Materials Research Science and Engineering Center supported by NSF grant DMR-2011750. A.M.-C. acknowledges support from the Princeton Center for Theoretical Science and the Human Frontier Science Program through the grant LT000035/2021-C

## VI. AUTHOR CONTRIBUTIONS

D.B.A. and S.S.D. designed the model; D.B.A. and S.S.D. designed the numerical simulations and theoretical analysis; B.A. performed all numerical simulations and theoretical analysis; D.B.A., A.M., and S.S.D. analyzed the data; S.S.D. designed and supervised the overall project. D.B.A. and S.S.D. discussed the results and implications and wrote the manuscript.eaba0353 (2020).

## SUPPLEMENTAL MATERIALS

## Notes

### Competing Interest Statement

The authors have declared no competing interest.

